# A broadly active flavivirus NS4B targeting compound that blocks NS4B-NS3 binding and replication organelle formation

**DOI:** 10.64898/2026.10.29.762806

**Authors:** Jessica L Smith, Theresa H. Nguyen, Sixue Zhang, Omar Moukha-Chafiq, Corinne E. Augelli-Szafran, Alec J. Hirsch

## Abstract

Mosquito-borne orthoflaviviruses, including West Nile virus (WNV), dengue virus (DENV), Zika virus (ZIKV), and Japanese encephalitis virus (JEV), cause substantial global morbidity and mortality, yet no broadly effective direct-acting antivirals are available. The viral nonstructural protein NS4B has emerged as a promising antiviral target, although the compounds described to date are largely DENV-specific. Here we characterize SRI-38841, a xanthine analog derived from hit-to-lead optimization of a previously described WNV inhibitor, which inhibits WNV replication with an IC₉₀ of 0.87 µM and low cytotoxicity, and retains activity against DENV, ZIKV, and JEV. Serial passage of WNV under compound selection yielded resistant viruses bearing mutations in NS4B, and reverse genetics was used to confirm the role of individual amino acid substitutions. Several of these residues map to regions analogous to those implicated in resistance to the DENV NS4B inhibitors JNJ-A07 and NITD-688. Docking of SRI-38841 to an AlphaFold2 model of WNV NS4B predicted a binding pocket within a cytosolic-facing funnel, and mutation of the pocket residues A45 and K56 likewise conferred resistance, while changes at Q170 and T220 did not. Together, these data favor a topology model in which the compound engages a cytosolic-facing pocket over one placing these residues in the ER lumen. Mechanistically, SRI-38841 disrupts the NS4B–NS3 interaction and abolishes replication organelle formation in infected cells. Unlike JNJ-A07 and NITD-688, which were inactive against WNV and ZIKV, SRI-38841 inhibits all four viruses tested, identifying this xanthine class as a starting point for broadly active orthoflavivirus antivirals.

**Importance:** Mosquito-borne orthoflaviviruses such as West Nile virus (WNV), dengue virus (DENV), and Zika virus (ZIKV) remain a substantial global health burden, and no broadly effective direct-acting antivirals are available. NS4B has emerged as one of the most promising targets in this space, but the inhibitors described to date, including the clinical-stage compounds JNJ-A07/JNJ-1802 and NITD-688, are largely DENV-specific. An antiviral with genuine pan-orthoflavivirus activity has been lacking. In this study we characterize SRI-38841, a xanthine analog arising from hit-to-lead optimization of a WNV inhibitor we previously described. SRI-38841 has broad spectrum activity, showing efficacy against DENV2, ZIKV, and JEV as well as WNV. It targets the NS4B protein, and and the resistance landscape converges with that of the aforementioned DENV inhibitors. Finally, docking of SRI-38841 against an AlphaFold2 model predicted a binding pocket within a cytosolic-facing funnel. These results favor the NMR/computational topology model over one placing these residues in the ER lumen — a distinction with direct consequences for how this inhibitor class is understood and optimized.

## Introduction

Mosquito-borne orthoflaviviruses, which include West Nile virus (WNV), Yellow fever (YFV), Japanese encephalitis (JEV), and the dengue viruses (DENV) among others, cause significant morbidity and mortality worldwide, with the majority of infections occurring in tropical and subtropical regions^1^. With the migration and expansion of the arthropod vectors that transmit these pathogens comes increased risk for these viruses to emerge in naïve populations^2^. Decades of research focused on the discovery of flavivirus-targeting antivirals have yielded only a handful of promising lead compounds, with most drugs failing in pre-clinical and early trials^3^. The search for effective antivirals that target these important human pathogens remains a critical pressing need.

The orthoflaviviruses are small, positive-stranded RNA viruses with a genome of approximately 10-11kb, made up of a single open reading frame (ORF), flanked by 5’- and 3’-untranslated regions (UTRs) that are essential for viral replication. The structural proteins (capsid (C), preMembrane (prM), and envelope (E)) are encoded at the 5’ end of the genome, followed by the nonstructural proteins (NS1, NS2A, NS2B, NS3, NS4A, NS4B, and NS5). The genome is translated as a single polyprotein that is cleaved co- and post-translationally by host and viral proteases^4^. Efforts to develop direct-acting antivirals (DAAs) against the flaviviruses are often focused on entry inhibitors (i.e., targeting C or E) or on the enzymatic factors, such as the viral protease, NS3, and the RNA dependent RNA polymerase (RdRp), NS5. However, in recent years, multiple compounds targeting the orthoflavivirus NS4B protein have been described, some of which have progressed to clinical trials (i.e., JNJ-1802/Mosnodenvir, NITD-688/EYU688). These and others have highlighted the viral protein NS4B as a promising target for DAA investigation^5–7^.

NS4B is a small multi-pass transmembrane protein that lacks enzymatic activity. Through direct interaction with other nonstructural proteins, it is reported to play an essential role in endoplasmic reticulum (ER) membrane remodeling and formation of replication complexes within the membrane invaginations termed replication organelles (ROs). In addition to its function as a scaffolding factor supporting the maintenance of replication complexes, NS4B also acts as a cofactor for NS3 helicase activity and promotes evasion of host innate responses by inhibiting interferon signaling^8,9^.

Recently we reported the discovery of a class of WNV-targeting antivirals and subsequent medicinal chemistry efforts to generate a lead molecule with reasonable pharmacokinetic properties^10^. In the present study, we describe the further characterization of the activity profile and mechanism of action of the lead molecule, SRI-38841 (previously, Compound 21). Interestingly, although SRI-38841 does not appear to be chemically similar to previously described flavivirus NS4B-targeting molecules^5–7^, we found that this compound directly targets WNV NS4B and exerts its antiviral effects through a similar mechanism.

## Results

### Identification of a class of broadly anti-orthoflaviviral compounds

Previous research from our group described the identification and detailed structure-activity relationship (SAR) study of a class of xanthine inhibitors that target WNV replication^10^. The hit compound, SRI-37776 (previously, Compound 1; Figure 1A, left) exhibited an inhibitory concentration (IC_90_) of 0.93 µM and reduced viral load (VLR) by >3-logs, with moderate cytotoxicity (CC_50_ =12 µM). Hit-to-lead optimization efforts on SRI-37776 led to the identification of SRI-38841 (previously, Compound 21; Figure 1A, right), with comparable IC_90_ and VLR values (IC_90_ = 0.87 µM, VLR = 3.5) and decreased toxicity (CC_50_ > 30 µM), as well as improved metabolic stability in mouse microsomes (t_1/2_ = 41 min) versus SRI-37776 (t_1/2_ = 2 min) (su). Dose response curves demonstrated that, in addition to its effect on WNV replication, SRI-38841 effectively inhibits replication of the related orthoflaviviruses, DENV2, ZIKV, and JEV (Figure 1B-E, Table 2A). These results highlight this class of xanthine molecules as attractive drug candidates with broad-spectrum anti-flavivirus activity.

**Figure 1.**
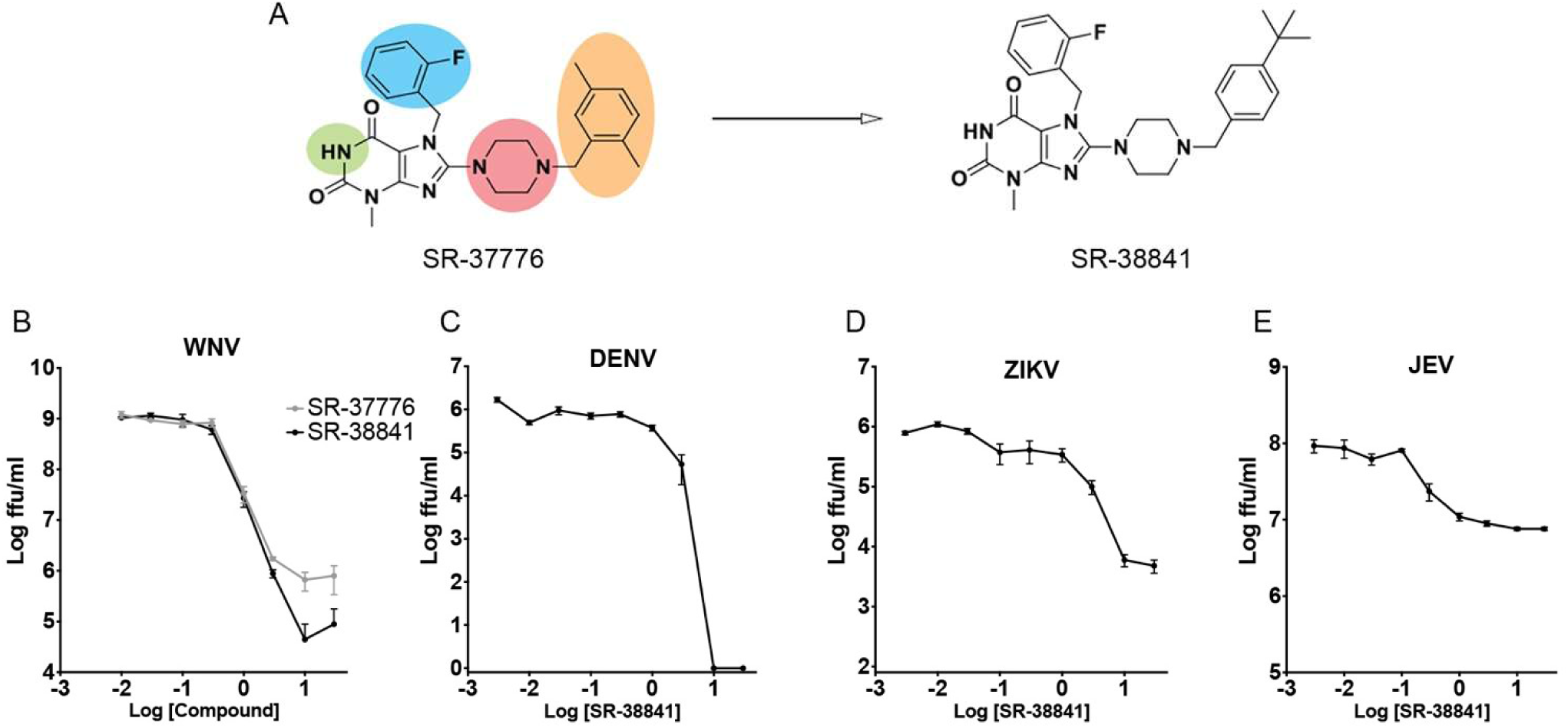
Identification of SRI-38841, a broadly acting anti-flaviviral small molecule. (A) Hit compound SRI-37776 (left), identified in a high throughput screen for WNV-targeting compounds. SAR analysis focused on modification of highlighted regions identified the lead compound, SRI-38841 (right), with more favorable pharmokinetic properties. HEK293 cells were infected in triplicate with WNV (B), DENV-2 (C), ZIKV (D), and JEV (E) at MOI=0.1 ffu/cell in the presence of 3-fold dilutions of compound (30µM→→0.041µM). At 48h (WNV, JEV) or 72h (DENV, ZIKV) p.i., supernatants were collected and infectious virus was quantified by focus forming assay on Vero cells.

**Table 1:** Efficacy and physical characteristics of SRI-37776 and SRI-38841.

|  | IC90 (uM) | IC50 (uM) | CC50 (uM) | VLR | Solubility (uM) | LogD | MLM t<sub>1/2</sub> (min) |
| --- | --- | --- | --- | --- | --- | --- | --- |
| **SRI-37776** | 0.93 ± 0.36 | 0.13 ± 0.01 | 12 | 3.2 | 9.7 | >4 | 1.6 |
| **SRI-38841** | 0.87 ± 0.02 | 0.11 ± 0.02 | >30 | 3.5 | 2.3 | >3.1 | 40.8 |

**Table 2:**
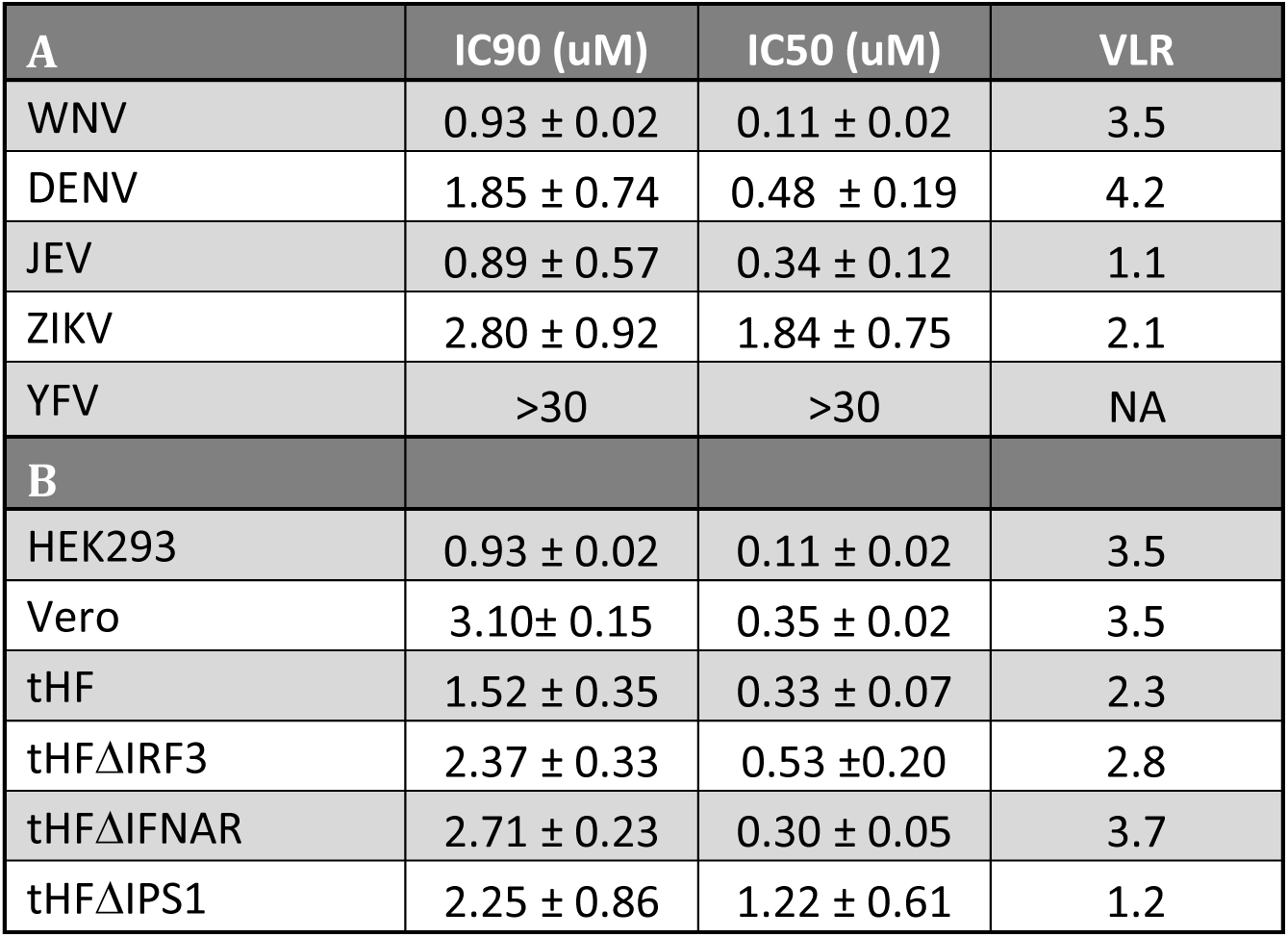
Efficacy of SRI-38841 against orthoflaviviruses. A. IC_50_ and IC_90_ values of SR-38841 against indicated orthoflavivirus. **B.** IC_50_ and IC_90_ values of SR-38841 against WNV in indicated cell type.

| **A** | IC90 (uM) | IC50 (uM) | VLR |
| --- | --- | --- | --- |
| WNV | 0.93 ± 0.02 | 0.11 ± 0.02 | 3.5 |
| DENV | 1.85 ± 0.74 | 0.48 ± 0.19 | 4.2 |
| JEV | 0.89 ± 0.57 | 0.34 ± 0.12 | 1.1 |
| ZIKV | 2.80 ± 0.92 | 1.84 ± 0.75 | 2.1 |
| YFV | >30 | >30 | NA |
| **B** |  |  |  |
| HEK293 | 0.93 ± 0.02 | 0.11 ± 0.02 | 3.5 |
| Vero | 3.10 ± 0.15 | 0.35 ± 0.02 | 3.5 |
| tHF | 1.52 ± 0.35 | 0.33 ± 0.07 | 2.3 |
| tHFΔIRF3 | 2.37 ± 0.33 | 0.53 ± 0.20 | 2.8 |
| tHFΔIFNAR | 2.71 ± 0.23 | 0.30 ± 0.05 | 3.7 |
| tHFΔIPS1 | 2.25 ± 0.86 | 1.22 ± 0.61 | 1.2 |

### SRI-38841 blocks WNV post-entry at an early step during replication

To investigate the mechanism of SRI-38841 antiviral activity, we measured the production of viral RNA (vRNA) using RT-qPCR in a single step growth curve (Figure 2A). SRI-38841 treatment (black bars) resulted in the reduction of vRNA by 1-2 logs at 16-24h post-infection, indicating that the compound is acting at a relatively early step in the life cycle, inhibiting the production of nascent viral genomes. To further explore the mechanism of action, we examined the inhibitory activity against WNV in several other cell types, including Veros, life extended (telomerase+) human fibroblasts (tHFs), and tHFs deficient in components of the type I interferon signaling system (tHF-ΔIRF3, -ΔIFNAR, -ΔIPS1). While replication levels varied among the cell lines, SRI-38841 retained comparable activity across the cell types tested, with IC_90_ values ranging from 0.9 µM to 3 µM (Figure 2B; Table 2). These data indicate that the inhibitory effects of SRI-38841 on WNV replication are not restricted to a specific cell type and are not strictly dependent on activation of interferon signaling.

**Figure 2.**
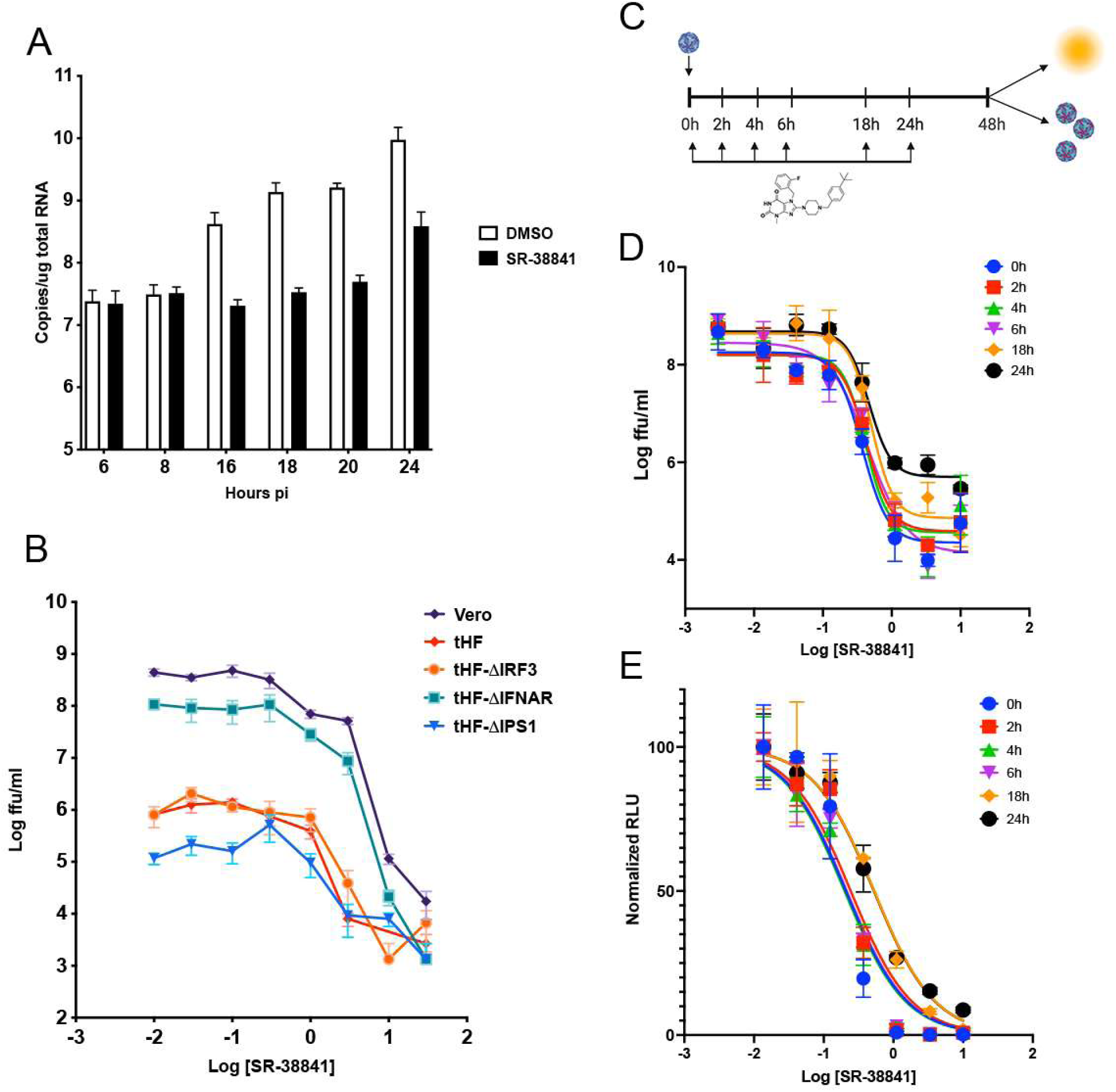
SRI-38841 inhibits viral replication at a step post-entry and is independent of type I interferon signaling. (A) HEK293 cells were infected at MOI=3ffu/cell. At indicated times p.i., total cellular RNA was collected and genomic RNA was quantitated by RT-qPCR. (B) Vero cells, telomerized human fibroblasts (WT), and fibroblasts knocked out for type I IFN signaling components (IRF3, IFNAR, IPS1) were infected with WNV at MOI=0.1ffu/cell in the presence of SRI-38841. Supernatants collected at 48h p.i. were quantitated by ffa. (C) Experimental procedure for time-of-addition experiments. HEK293 cells were infected with WNVnLuc at MOI=0.1 ffu/cell and at indicated times p.i., 3-fold dilutions (30µM→→0.041µM) of SRI-38841 was added. (D) Supernatants collected at 48h p.i. were assayed for infectious virus by ffa. (E) Luciferase activity of cells was quantitated at 48h p.i. using the NanoGlo luciferase assay kit.

To investigate the step during the viral life cycle at which SRI-38841 is exerting its effects, we performed time of addition experiments, with the reasoning that retention of activity would signify a post-entry mechanism of inhibition when SRI-38841 was added at later timepoints. Low MOI (0.5 ffu/cell) infections were performed in HEK293 cells with a WNV reporter virus expressing nanoluciferase (nLuc), and at indicated times post-infection (0-24h), media was removed from cells and replaced with a series of concentrations of SRI-38841 (10mM→10nM). At 48h p.i., supernatants were removed and assayed for infectious virus by a focus forming assay. Concurrently, nLuc expression was quantified in cell lysates (Figure 2C). Compound addition as late as 6h p.i. retained equivalent inhibitory effects on nLuc expression (Figure 2D; Table 3) and infectious virus production (Figure 2E; Table 3). Compound addition at later timepoints (18h, 24h) resulted in slightly reduced activity in both assays; however, this reduction was inconsistent with a mechanism by which SRI-38841 is acting on viral entry. We also performed these assays in a range of other cell types with comparable results (Table 3), although the loss of activity against nLuc expression observed at later timepoints was more pronounced in Huh7.5 and Vero cells. These results, coupled with the RT-qPCR findings (Figure 2A), indicate that the antiviral effects exerted by SRI-38841 are occurring post-entry, and the mechanism likely involves direct inhibition of genome replication.

**Table 3:** Inhibition of WNV replication by SRI-38841 added up to 24h post infection. Concentration of SR-38841 resulting in 50% reduction of nLuc activity from an nLuc-expressing WNV (IC_50_), or a 10-fold reduction in infectious progeny (IC_90_) shown for indicated cell types at indicated times of compound addition.

| **Table 3** |  |  |  |  |  |  |  |  |
| --- | --- | --- | --- | --- | --- | --- | --- | --- |
| Time of compound addition | nLuc IC50 (uM) |  |  |  | IC90 (uM) |  |  |  |
|  | HEK293 | Huh7 | Huh7.5 | Vero | HEK293 | Huh7 | Huh7.5 | Vero |
| **0h** | 0.22 ± 0.06 | 0.59 ± 0.14 | 1.58 ± 0.06 | 2.39 ± 0.20 | 1.01 ± 0.13 | 2.56 ± 0.34 | 2.71 ± 0.04 | 2.53 ± 0.14 |
| **2h** | 0.24 ± 0.02 | 0.67 ± 0.11 | 1.80 ± 0.12 | 3.21 ± 0.11 | 1.25 ± 0.49 | 2.18 ± 0.03 | 4.42 ± 0.14 | 3.12 ± 0.18 |
| **4h** | 0.20 ± 0.01 | 0.71 ± 0.13 | 1.92 ± 0.10 | 3.12 ± 0.10 | 0.95 ± 0.19 | 2.11 ± 0.14 | 5.71 ± 0.97 | 3.35 ± 0.58 |
| **6h** | 0.21 ± 0.02 | 0.89 ± 0.25 | 1.81 ± 0.27 | 4.19 ± 0.53 | 1.67 ± 0.78 | 2.16 ± 0.09 | 5.43 ± 1.76 | 4.16 ± 0.54 |
| **18h** | 0.54 ± 0.10 | 1.96 ± 0.18 | 3.39 ± 0.55 | 3.37 ± 0.12 | 2.37 ± 0.07 | 3.27 ± 0.015 | 7.76 ± 0.29 | 3.33 ± 0.02 |
| **24h** | 0.55 ± 0.19 | 2.39 ± 0.69 | 6.18 ± 0.78 | 11.16 ± 2.48 | 3.06 ± 0.08 | 4.70 ± 0.32 | 7.84 ± 0.41 | 3.79 ± 0.48 |

### Identification of WNV NS4B as a direct target of SRI-38841

To determine whether the xanthine analogs are functioning as direct-acting antivirals (DAAs) targeting a specific viral factor, we passaged virus in the presence of compound to induce the establishment of compound resistant viruses. HEK293 cells were infected in the presence of 1mM SRI-37776 and passaged 8 times. Virus from each passage was evaluated for its sensitivity to SRI-37776 inhibition (Figure 3A). As expected, wild type and control (0.1% DMSO) passaged virus was highly susceptible to treatment with 10 µM SRI-37776, resulting in >2-log reduction in viral titer. In contrast, SRI-37776-passaged virus quickly lost sensitivity and demonstrated complete resistance after only four passages. Five isolates independently passaged 8 times were deep sequenced, and clusters of mutations were identified that mapped to the nonstructural protein, NS4B (Figure 3B). Some mutations, for example V88A and V173A, emerged in several of the independently passaged isolates. To determine the role that each of the identified mutations play in sensitivity to compound treatment, we introduced individual residue changes into a WNV infectious clone^11^ and measured the effect of these mutations on SRI-38841 inhibition. Introduction of the single NS4B mutations, V88A and S93P, conferred resistance to compound treatment, resulting in a shift in IC_90_ values from 900nM to >10µM (Figure 3C; Table 4). Conversely, incorporation of the changes at T115I or V173A did not affect the sensitivity of these viruses to SRI-38841. Notably, the changes at T115 and V173 were only observed to occur concurrently with confirmed resistance-conferring mutations (V88A and S93P). Attempts to recover viruses mutated at A113, Y122, and I229 were unsuccessful, possibly due to a negative impact these mutations might have on viral replication. We also investigated the effect of mutating the resistant sites, V88 and S93, to other residues with variable results. While mutation of the valine at position 88 to a serine (the residue at the equivalent position of DENV NS4B) also produced resistant virus, mutation of the serine at position 93 to either alanine or glycine had no effect (Figure 3C; Table 4). These results indicate that the xanthine analogs, SRI-37776 and SRI-38841, are DAAs targeting WNV NS4B and identify a putative region where the compounds are directly interacting with the viral protein.

**Figure 3.**
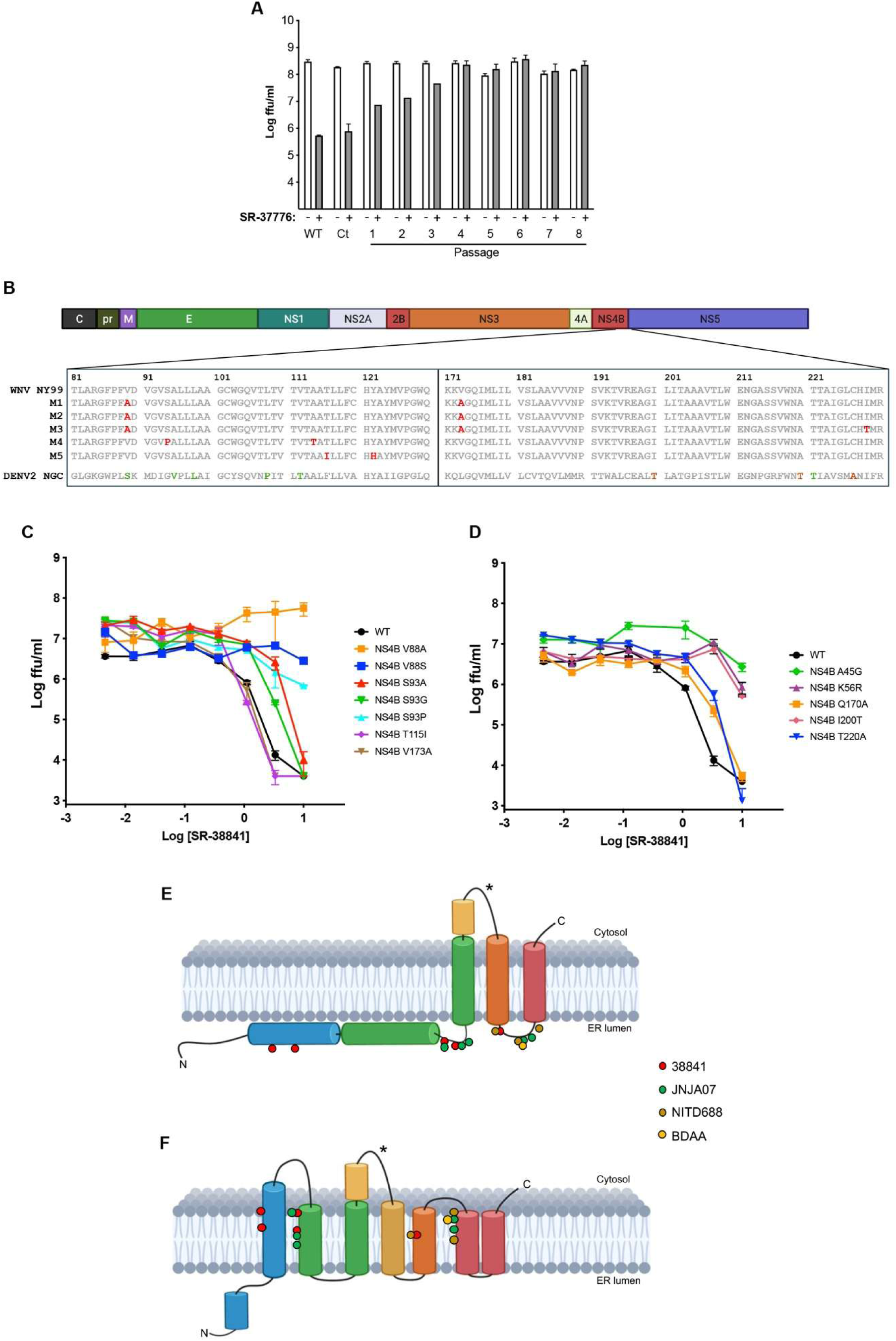
Identification of WNV NS4B as the target of xanthine compounds. (A) WNV was passaged on HEK293 cells in the presence of 1µM SRI-37776 or DMSO (Ct). Virus collected at day 3 for each passage was assayed for its sensitivity to 10µM SRI-37776 (grey bars). (B) Region of WNV NS4B where clusters of mutations were identified in independently generated SRI-37776-resistant WNV mutants (M1-M5; passage 8) and the corresponding region of DENV2 NS4B with previously identified mutations that confer resistance to JNJ-A07 (green) and NITD-688 (brown) (C-D) NS4B mutations were introduced into a WNV infectious clone. HEK293 cells were infected with WT and mutant viruses in the presence of 3-fold dilutions of SRI-38841 and supernatants assayed by focus-forming assay. (E-F) Divergent models of flavivirus NS4B structure. (E) Model predicted by biochemical methods predicts first 110 residues and all identified SRI-38841-resistance conferring mutations reside in endoplasmic reticulum lumen. (F) Model predicted through a combined Alphafold/NMR approach proposes that NS4B spans ER membrane 7 times, forming a funnel structure (Supplemental Figure 1) that faces the cytosol. SRI-38841-resistance conferring mutations reside within the funnel in this model. Circles indicate positions of resistance-conferring residues (red: SRI-38841; green: JNJ-A07; brown: NITD-688; yellow: BDAA). Star: NS4B/NS3 interaction site.

**Table 4:** Calculated IC_90_ values for SRI-38841 of WNV with amino acid substitutions in NS4B. Highlighted mutants indicate complete resistance. *Mutants identified by serial passage in the presence of SRI-38841.

|  | IC<sub>90</sub> (uM) |
| --- | --- |
| WNV | 0.84 ± 0.19 |
| WNV NS4B W42A | 3.24 ± 0.19 |
| WNV NS4B A45G | >10 |
| WNV NS4B K56R | >10 |
| WNV NS4B V88A* | >10 |
| WNV NS4B V88S | >10 |
| WNV NS4B D89A | 2.07 ± 0.38 |
| WNV NS4B G91D | 2.20 ± 0.15 |
| WNV NS4B V92A | 1.62 ± 0.42 |
| WNV NS4B S93A | 3.05 ± 0.8 |
| WNV NS4B S93G | 1.70 ± 0.35 |
| WNV NS4B S93P* | >10 |
| WNV NS4B L95A | 2.23 ± 0.39 |
| WNV NS4B L96A | 3.29 ± 0.26 |
| WNV NS4B T115I | 0.97 ± 0.31 |
| WNV NS4B Q170A | 2.47 ± 0.08 |
| WNV NS4B K171A | 2.93 ± 0.44 |
| WNV NS4B V173A* | 0.71 ± 0.16 |
| WNV NS4B Q175A | 3.03 ± 0.25 |
| WNV NS4B I200T | >10 |
| WNV NS4B A220T | 1.67 ± 0.58 |
| WNV NS4B T221A | 2.39 ± 0.41 |
| WNV NS4B T222A | 2.10 ± 0.11 |
| WNV NS4B W242F | 2.66 ± 0.30 |
| WNV NS4B K250N | 3.10 ± 0.33 |

Previous antiviral drug discovery efforts have identified multiple classes of flavivirus targeting small molecules (JNJ-A07, NITD-688) that exert their antiviral effects through direct interaction with DENV NS4B^5,6,12–14^. Interestingly, many of the resistance-conferring mutations described for these DENV NS4B targeting compounds map to analogous regions of WNV NS4B identified in this study (Figure 3B, bottom; JNJ-A07/-1802 [green] ^5,14^, NITD-688 [brown] ^6,12^). Analysis of the structural similarities shared among these three classes indicates that each contain a bicyclic core of a 6-membered ring fused with a 5-membered ring (namely, xanthine core of SRI-38841, tetrahydrothienopyridine core of NITD-688, and indoline core of JNJ-A07; Supplemental Figure 2), and all three classes of compounds are likely exerting antiviral effects through similar mechanisms. To explore further the analogous activities of these compounds, we analyzed the effects of changing residues at and around these sites of susceptibility of WNV to SRI-38841 (Table 2). Mutation of the isoleucine at position 200 of WNV NS4B (equivalent to DENV NS4B T195, a resistance site for NITD-688) conferred resistance to SRI-38841 (Figure 3D), providing further evidence that the inhibitory mechanism of SRI-38841 is parallel to that of the DENV targeting compounds, JNJ-A07 and NITD-688.

Although the crystal structure of flavivirus NS4B has not been solved, multiple structural models have been proposed based on biochemical studies^15^, nuclear magnetic resonance (NMR)^16,17^, and computational modeling^9^ (Figure 3E and 3F). Five transmembrane domains (TMD) are predicted based on hydrophobicity and helical structure; however, the various models differ in their prediction of whether the first two putative TMDs (pTMD1 and pTMD2) cross the membrane or are located within the ER lumen. The model built from biochemical data (Figure 3E) predicts that the resistance-conferring mutations described for these classes of compounds are present in the ER lumen. Alternatively, the structure predicted from NMR and computational modeling (Figure 3F) proposes these residues lie within a funnel-like structure facing the cytosol (Supplemental Figure 1). Docking studies of SRI-38841 with an Alphafold2 generated structure of WNV NS4B predicts residues (Suppl Figure 1B) that form a putative binding pocket for SRI-38841 within the funnel structure have a predicted docking affinity of -10.9 kcal/mol^18,19^. Critical interactions between residues of the binding pocket and the ligand (Suppl Figure 1C) were predicted (hydrophobic interactions: grey, hydrogen bonds: blue) using the Protein-Ligand Interaction Profiler (PLIP; Suppl Tables 1, 2)^20^. We explored the impact of mutating residues within this putative binding pocket and found that, while mutation of A45 and K56 resulted in resistance, changes at Q170 and T220 did not affect susceptibility to SRI-38841 treatment (Figure 3D; Table 4). In conjunction with the resistance data presented above, these results are supportive of the NMR/computationally predicted topology model of NS4B structure (Figure 3F) in which SRI-38841 binds directly within a cytosolic facing binding pocket (Suppl Figure 1C).

Orthoflavivirus NS4B is a transmembrane protein with no known enzymatic activity that is essential for viral replication. Translation of the flavivirus polyprotein occurs at the ER, and cleavage of NS4A-2K-NS4B by host and viral proteases initiates ER membrane remodeling, leading to the formation of ER-derived membranous structures, including convoluted membranes (CMs) and vesicle packets (VPs)^21,22^. Replication of the viral genome occurs at replication complexes located within VPs, also called replication organelles (ROs). Within the ROs, NS4A and NS4B act as scaffolds and recruit other NS proteins to support the formation of replication complexes. In addition to the scaffolding function of NS4B, it has also been demonstrated to be a cofactor for NS3 helicase activity during viral genome replication^8,9^. The NS4B-NS3 interaction has been shown to be disrupted by the DENV NS4B targeting compounds, JNJ-A07 and NITD-688^12,14^. To investigate whether SRI-38841 acts against WNV replication through a similar mechanism, we performed co-immunoprecipitations with Flag-tagged WNV NS4B and myc-tagged WNV NS2B/3 (Figure 4A). As expected, immunoprecipitation of Flag-WNV NS4B efficiently pulled down WNV NS3; however, treatment with 1µM or 10µM SRI-38841 abolished this interaction. Additionally, we found that introduction of the resistance conferring mutations, V88A and S93P, did not greatly impact the NS4B/NS3 interaction but the susceptibility of this interaction to SRI-38841 treatment was lost with these mutants. These data demonstrate that SRI-38841 inhibits WNV replication by blocking the NS4B/NS3 interaction, in a similar mechanism to the DENV NS4B targeting compounds. Interestingly, the putative binding region identified in this study and in previous studies is in a distinct area of NS4B than that defined for the NS4B/NS3 interaction (Figure 3E, stars)^23^. This suggests that these small molecules are not acting by directly obstructing NS4B/NS3 binding but are more likely altering the conformation of NS4B, resulting in the disruption of this interaction.

**Figure 4.**
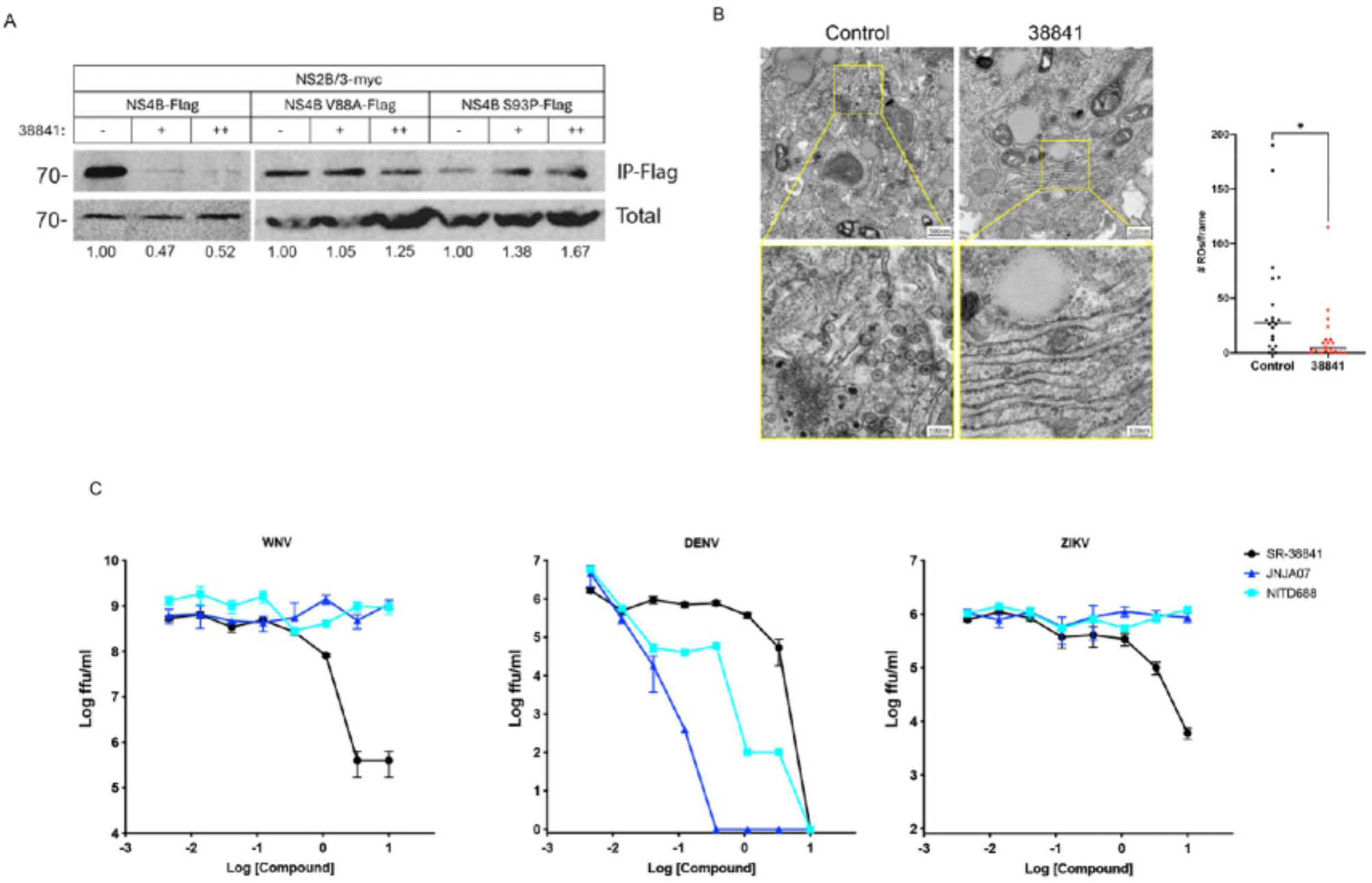
Xanthine compounds target WNV NS4B-NS2B/3 interaction. (A) HEK293 cells were co-transfected with WNV NS4B-Flag (wild-type, V88A, or S93P) and WNV NS2B/3-myc. At 6h post-transfection, media was changed and replaced with DMSO (-), 1µM (+), or 10µM (++) SRI-38841. At 48h p.t., cells were lysed and immunoprecipitated with anti-Flag sepharose beads. Immunoprecipitated and total lysate fractions were assayed by western blot for myc tag. (B) Huh7 cells were infected with WNV at MOI=0.5 ffu/cell +DMSO or 10µM SRI-38841 for 24h before fixation and eletron microscopy analysis. 20 frames per sample were captured at ___X. Frames were randomly assigned and the total number of replication organelles (ROs)/frame were blindly counted. (C) HEK293 cells were infected in triplicate with WNV, DENV, and ZIKV at MOI=0.1 ffu/cell in the presence of 3-fold dilutions of compound (30µM→→ 0.041µM). At 48h (WNV) or 72h (DENV, ZIKV) p.i., supernatants were collected and infectious virus was quantified by focus forming assay on Vero cells.

In addition to the disruption of NS4B and NS3 protein-protein interaction, the DENV NS4B-targeting compounds, JNJ-A07 and NITD-688, have been demonstrated to block the formation of ROs^14^. To determine whether RO formation in WNV-infected cells is similarly affected by treatment with SRI-38841, WNV-infected cells treated with DMSO (control) or 5µM SRI-38841 were fixed at 24h and analyzed by electron microscopy. To quantify ROs in each sample, 20 random fields were blinded and the number of ROs/frame counted. As expected, ROs were readily observed in control-treated infected cells, while treatment with SRI-38841 significantly reduced the number to mostly undetectable levels (Figure 4B). These results further support the theory that all of these NS4B-targeting compounds are acting by a similar mechanism to block flavivirus infection.

Because of the parallel mechanism observed for the DENV and WNV NS4B-targeting compounds, we sought to understand the breadth of activity of each of these classes of small molecules against the flaviviruses. Dose-response curves were performed with SRI-38841, JNJ-A07, and NITD-688 against DENV, WNV, and ZIKV (Figure 4C; Table 5). Interestingly, we found that SRI-38841 was broadly effective against the three flaviviruses tested, with IC_90_s in the single micromolar range. While SRI-38841 displayed lower efficacy against DENV replication than both JNJ-A07 and NITD-688, neither of the DENV-targeting compounds demonstrated any inhibitory activity against WNV or ZIKV. These data suggest that, although SRI-38841 is a less robust inhibitor of DENV replication, its activity extends to related flaviviruses, making this class of compounds an attractive target for development as broadly active DAAs.

**Table 5:** Efficacy of SRI-38841 compared to previously described NS4B targeting anti-DENV compounds.

|  | IC90 (uM) |  |  |
| --- | --- | --- | --- |
|  | WNV | DENV2 | ZIKV |
| SRI-38841 | 1.06 ± 0.04 | 1.60 ± 0.26 | 3.19 ± 0.16 |
| JNJA07 | >10 | 0.014 ± 0.002 | >10 |
| NITD688 | >10 | 0.019 ± 0.002 | >10 |

## Discussion

In this study we present the characterization of a class of xanthine compounds with broad anti-flavivirus activity that exert inhibitory effects through direct interaction with the viral NS4B protein. We demonstrate that, although the structure of SRI-38841 differs considerably from that of previously identified flavivirus NS4B inhibitors, such as the DENV-specific, JNJ-A07 and NITD-688^5,10,12^, all of these compounds appear to be blocking viral replication through parallel mechanisms. Through direct interaction with NS4B, these compounds exert antiviral effects via disruption of NS4B/NS3 binding and inhibition of RO formation, wherein viral replication is occurring^13,14^. The repeated emergence of NS4B-targeting small molecules from antiviral drug discovery programs highlights the vital importance of this viral factor in the life cycle and its substantial potential as a drug target for future DAA screening efforts.

Because of the highly membranous nature of flavivirus polyprotein translation and genome replication sites, a comprehensive understanding of nonstructural viral protein functions and structures remains elusive. As the tertiary structure of NS4B has not been resolved, uncertainty remains about its exact conformation and topology, and competing models have been proposed (Figure 4A)^15–17,24^. The primary difference between these predicted models lies in whether the first ∼100 amino acids (pTM1 and pTM2) cross the membrane or are located within the ER lumen. Furthermore, it has been suggested that different flavivirus NS4Bs (i.e., DENV vs ZIKV) may have distinct topological structures^25^. Cleavage from NS4A and NS5 during co-translational processing, as well as interactions with host proteins, may also impact the NS4B structure^15,25–28^. When considering the resistance-conferring mutations described previously for DENV NS4B and for WNV NS4B which is reported in this study, the different models place the putative ligand-docking region either contained entirely in the ER lumen (biochemical model; Figure 4A, top) or within a funnel-like structure embedded in the ER membrane facing the cytosol (NMR/computational; Figure 4A, bottom; Suppl Figure 1). More research is needed to resolve these disparate theories and gain a better understanding of how NS4B-targeting antiviral drugs interact with and disrupt viral protein function.

Although traditional antiviral discovery efforts tend to emphasize compounds that target enzymatic viral factors, the independent identification by multiple groups of effective flavivirus-targeting drugs that function by acting on the nonenzymatic NS4B protein supports an alternative approach that focuses on targeting NS4B and other NS proteins within the replication complex. Furthermore, the multifunctional nature of the viral NS proteins makes them ideal candidates for therapeutic targeting. The work presented here further supports the development of flavivirus NS4B binding compounds as effective therapeutics and offers insight into expanding the breadth of these molecules to target a range of flaviviruses.

## Acknowledgements

This work was supported by grants from the National Institutes of Health 1U19AI109680-01 and 1U19AI142759-01. Electron microscopy was performed at the Multiscale Microscopy Core, a member of the OHSU University Shared Resource Cores, RRID: SCR_009969. Thermo Fisher Scientific Talos 120C was funded in part by the M.J. Murdock Charitable Trust. We would specifically like to acknowledge Rodolfo Alvarado who assisted with this project.

## Materials and Methods

### Cell culture and reagents

HEK293, C6/36, Huh7, tHF, and Vero cells were maintained in Dulbecco’s modified Eagle medium (DMEM; Gibco) supplemented with 10% fetal bovine serum (Hyclone), 2 mM l-glutamine (Invitrogen), 100 U/ml penicillin G sodium, 100 μg/ml streptomycin sulfate (Invitrogen). Telomerized human foreskin fibroblasts (tHFs) and tHF knockouts (DIRF3, D IFNAR1, D IPS1) were a gift from V. DeFilippis, Oregon Health & Science University (OHSU) ^29^.

### Virus strains

DENV2 (New Guinea C) was obtained from ATCC. West Nile virus (385–99) has been previously described^3^. Zika virus strain PRVABC59 was isolated by the Centers for Disease Control (CDC) from an individual in Puerto Rico in December 2015 ^30^. JEV (SA-14-2-8) was obtained from Robert Tesh, World Reference Center for Emerging Viruses and Arboviruses, University of Texas Medical Branch. Stocks were passaged twice on C6/36 cells. Supernatants were pelleted over a 20% sorbitol cushion and resuspended in DMEM/0.1%FBS. Stock titers were determined by focus forming assay, as previously described^31^. WNVnLuc was a kind gift of R. Baric (UNC Chapel Hill).

### Dose response curves and IC90 determination

Dose response curves were performed at MOI = 0.1 ffu/cell in the presence of 8x3-fold dilutions of compound in triplicate. At 48h pi, supernatants were collected and titrated by focus forming assay. IC90 values were determined by fitting normalized data points to a nonlinear regression model and calculating concentration at which viral titers were reduced by 90% (1-log) using GraphPad Prism software.

### Time-of-addition assay

HEK293 or Huh7 cells were infected with WNV or WNVnLuc at MOI = 0.1 ffu/cell. At indicated times p.i. media was removed and replaced with 8x3-fold dilutions of compound in triplicate. Supernatants were collected at 48h post-infection and quantitated by focus forming assay. Luciferase production was quantitated using the NanoGlo Luciferase Assay System (Promega #N1130) according to manufacturer’s instructions.

### RNA replication assay

HEK293 cells were infected with WNV at MOI = 3 ffu/cell +1µ**M** SRI-38841 or DMSO control. At indicated times post-infection, RNA was isolated from supernatants and analyzed by RT-qPCR using a primer/probe set designed to amplify a region within the NS3 gene of the WNV genome (Forward: 5′-GCACTGAGAGGACTGCCCAT-3’; Reverse: 5′-TGGGTGAGGGTAGCATGACA-3′; Probe 5′FAM-TACCAGACATCCGCAGTGCCCAGA-MGB-3′)^32^.

### Resistance mutant generation

HEK293 cells were infected at MOI = 0.1 ffu/cell in the presence of 1µM SRI-37776. Supernatants were collected at 48h p.i. and passaged onto naïve HEK293 cells + 1µM SRI-37776. Control viruses were passaged in 0.1% DMSO. Virus collected at each passage was assayed for resistance to inhibitor treatment and deep sequenced. Results are reflective of more than 5 independent derivations of resistant viruses.

### Infectious clone

Mutant viruses were produced from the WNV (NY99) infectious clone plasmids, pWN-AB and pWN-CG, as previously described^11^. Ligated template was in vitro transcribed using the HiScribe T7 ARCA mRNA in vitro transcription kit (NEB E2065). Ivt reactions were electroporated into 1 × 10^7 Vero cells, and media was changed the following day. Supernatants were collected when visible CPE was observed (5-8d post-transfection), centrifuged to remove cellular debris, and purified over a 20% sorbitol cushion. Dose-response curves were performed to determine susceptibility to SRI-38841 treatment, as above. Introduction of NS4B mutations into the pWN-CG plasmid was performed by site-directed mutagenesis using the QuikChange protocol with Phusion Taq mix (Thermofisher F530). Successful mutagenesis was confirmed by Sanger sequencing.

### Immunoprecipitations

HKE293 cells were transfected with plasmids expressing Flag-tagged WNV NS4B and myc-tagged WNV NS2B/3 in 6-well plates using lipofectamine 3000 (ThermoFisher 3000008), according to the manufacturer’s instructions. At 6h pt, media was removed and replaced with DMSO, 1µM, or 10µM SRI-38841. Lysates were collected at 48h pt in lysis buffer (10mM TrisHCl pH7.4, 150mM NaCl, 1% TNX100) and immunoprecipitations were performed using anti-Flag Affinity gel (Sigma A2220). Total lysates and immunoprecipitated samples were analyzed by SDS-PAGE/Western blot with an anti-myc antibody (Fisher R950-25).

### Computational modeling and ligand docking

The WNV (NY99; accession no MZ605381) and DENV2 (NGC; accession no MK506264) NS4B amino acid sequences were predicted with Alphafold 2 using the ColabFold interface^33,34^. Pdb files were analyzed and aligned in PyMol (v2.5.2). SRI-38841 binding to WNV NS4B was predicted with the AutoDock Vina (v1.2.0) PyMol plug-in^18,19^. Hydrophobic and hydrogen bond interactions were measured and visualized with Protein-Ligand Interface Profiler (PLIP)^20^.

### Electron microscopy

Huh7 cells were infected with WNV at MOI=0.5 ffu/cell + DMSO or 10µM SRI-38841. At 24h p.i., cells were scraped in PBS, pelleted, then fixed with 2.5% glutaraldehyde, 2% paraformaldehyde prepared in 0.1M sodium cacodylate buffer containing 0.1M sucrose (pH: 7.4). Pelleted cells were buffer washed then encapsulated in buffered 3% agarose. All subsequent steps were microwave-assisted with a laboratory microwave. Samples were washed in buffer, post-fixed in reduced osmium consisting of 2% aq. OsO4 and 1.5% aq. potassium ferricyanide, washed in water, stained in en bloc consisting of 1% aq. uranyl acetate, and dehydrated through a graded acetone series (50% to 100%), followed by 100% anhydrous acetone. Dehydrated samples were infiltrated with a 1:1 mixture of anhydrous acetone:Epon epoxy resin containing Z6040 embedding primer, followed by 100% resin. Samples were embedded in coffin molds using fresh resin and polymerized at 60°C for a minimum of 24 hours. Ultrathin sections (∼100 nm) were collected on 100-mesh copper grids coated with Formvar and carbon and counterstained with 2% aq. uranyl acetate and Reynold’s lead citrate. Sections were examined using a Thermo Fisher Scientific Talos L120C transmission electron microscope operated at 80 kV. TEM images were acquired with a CETA 16M camera using Velox software. 20 frames were captured per sample, and images were randomly sorted before replication organelle quantification.

## Supplemental figures

**Supplemental Figure 1.**
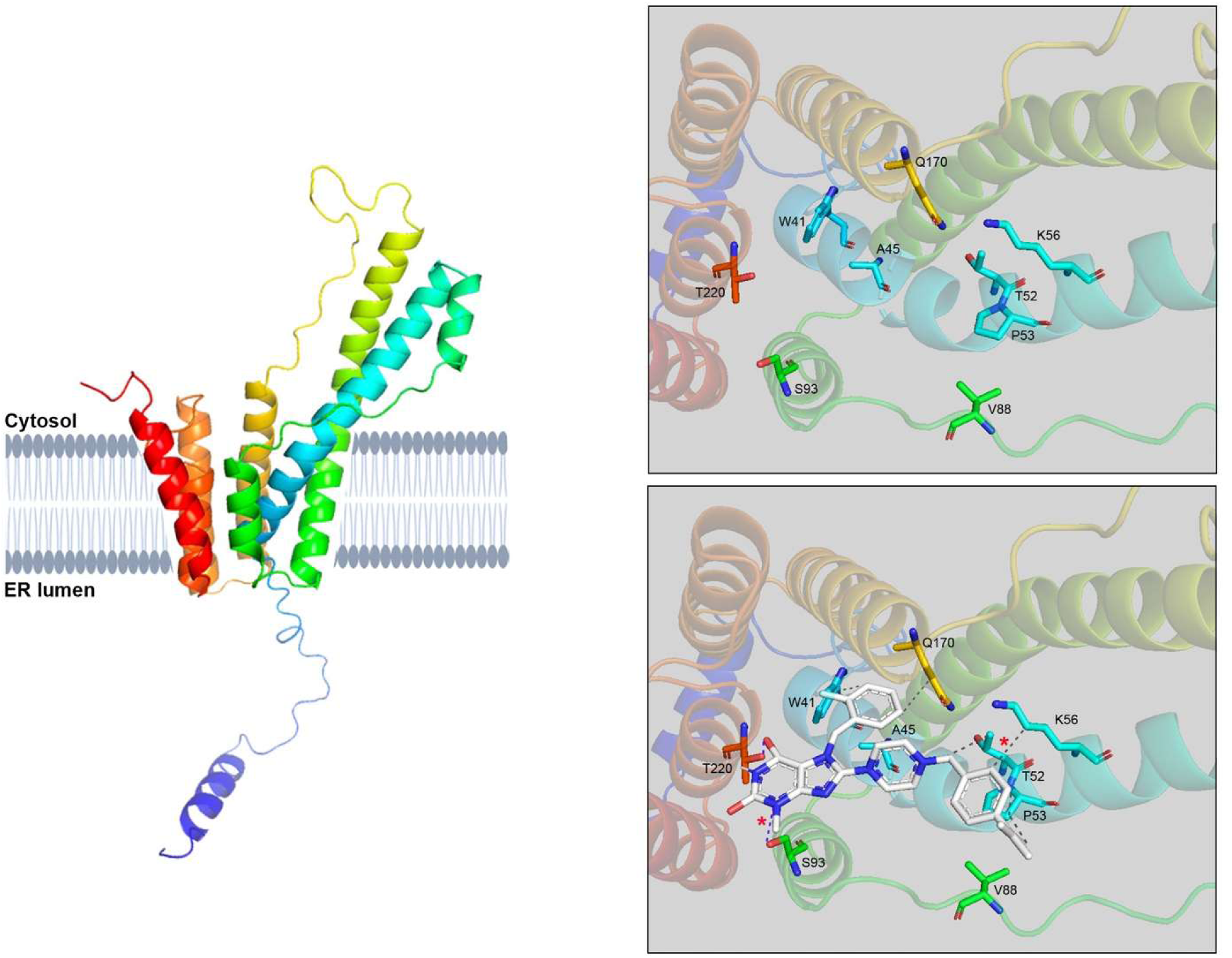
(A) Alphafold/NMR structure of WNV NS4B illustrating the cytosolic-facing funnel structure formed by the spanning of the ER membrane by 7 transmembrane domains. (B) Topdown view (from cytosol) of funnel structure highlighting residues within the funnel predicted to interact with SRI-38841 by AutoDock Vina (affinity= -10.9 kcal/mol) (C) Interactions predicted between WNV NS4B and SRI-38841: hydrophobic interactions (grey dashes), hydrogen bonds (blue dashes). Red stars indicate predicted interactions with resistance-conferring residues identified in this study.

**Supplemental Figure 2.**
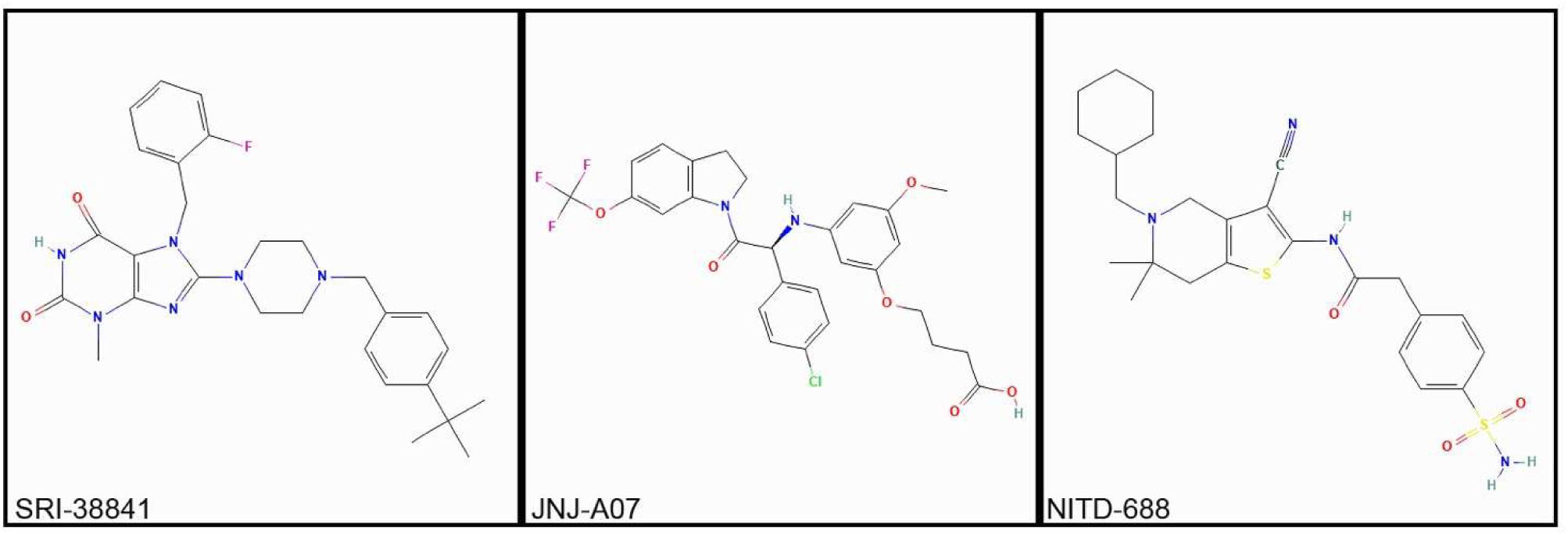
Molecular structures of three classes of flavivirus NS4B targeting compounds: SRI-38841 (PubChem CID: 155410050), JNJ-A07 (PubChem CID: 162468776), NITD-688 (PubChem CID: 146279852)

## Supplemental tables

**Supplemental table 1:**
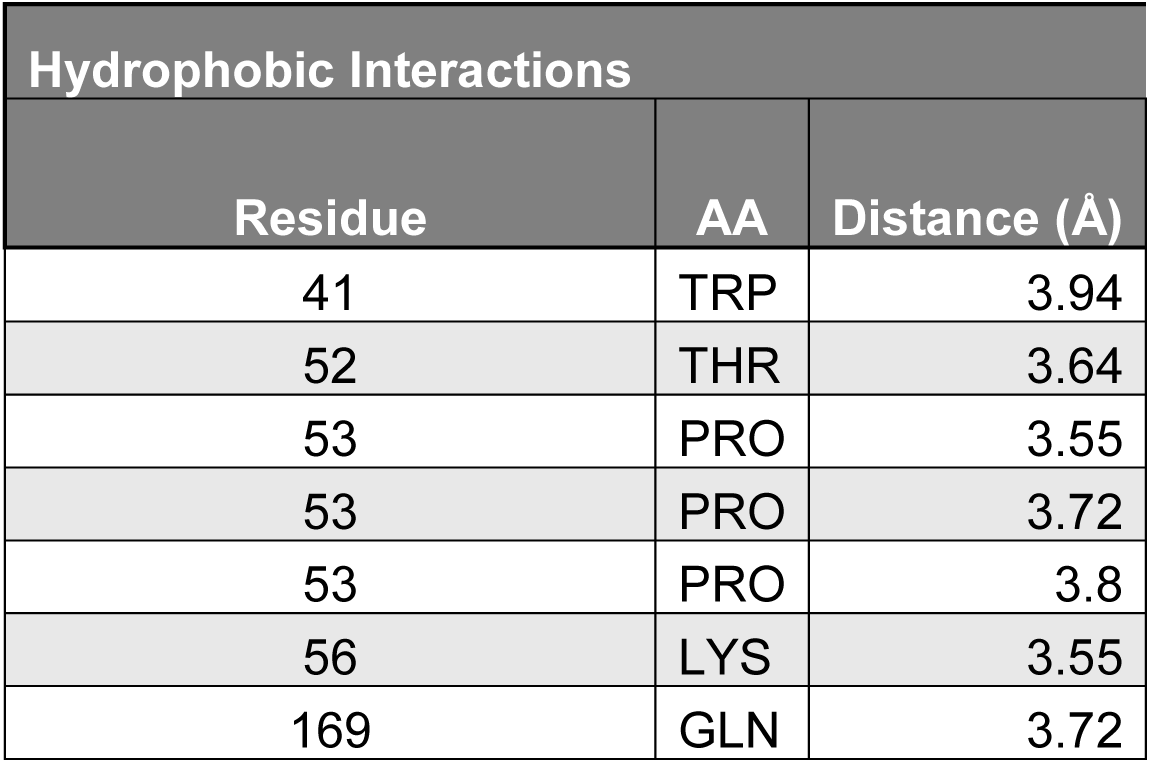
Predicted hydrophobic interactions between NS4B residues and SRI-38841. Interactions as shown in supplemental figure 1C (gray dashed lines).

**Supplemental table 2:** Predicted hydrogen bond interactions between NS4B residues and SRI-38841. Interactions as shown in supplemental figure 1C (blue dashed lines).

| Hydrogen Bonds |  |  |  |
| --- | --- | --- | --- |
| Residue | AA | Distance (Hydrogen-Acceptor) (Å) | Distance (Donor-Acceptor) (Å) |
| 93 | SER | 2.58 | 3.49 |
| 220 | THR | 2.16 | 2.72 |

## References

1. Pierson, T. C. & Diamond, M. S. The continued threat of emerging flaviviruses. Nat. Microbiol. 5, 796–812 (2020).

2. Weaver, S. C. Urbanization and geographic expansion of zoonotic arboviral diseases: mechanisms and potential strategies for prevention. Trends Microbiol. 21, 360–363 (2013).

3. Tripathi, A., Chauhan, S. & Khasa, R. A Comprehensive Review of the Development and Therapeutic Use of Antivirals in Flavivirus Infection. Viruses 17, (2025).

4. Chambers, T. J., Hahn, C. S., Galler, R. & Rice, C. M. Flavivirus genome organization, expression, and replication. Annu. Rev. Microbiol. 44, 649–688 (1990).

5. Kaptein, S. J. F. et al. A pan-serotype dengue virus inhibitor targeting the NS3-NS4B interaction. Nature 598, 504–509 (2021).

6. Moquin, S. A. et al. NITD-688, a pan-serotype inhibitor of the dengue virus NS4B protein, shows favorable pharmacokinetics and efficacy in preclinical animal models. Sci. Transl. Med. 13, eabb2181 (2021).

7. Guo, F. et al. A Novel Benzodiazepine Compound Inhibits Yellow Fever Virus Infection by Specifically Targeting NS4B Protein. J. Virol. 90, 10774–10788 (2016).

8. Zmurko, J., Neyts, J. & Dallmeier, K. Flaviviral NS4b, chameleon and jack-in-the-box roles in viral replication and pathogenesis, and a molecular target for antiviral intervention. Rev. Med. Virol. 25, 205–223 (2015).

9. Wang, Y., Xie, X. & Shi, P.-Y. Flavivirus NS4B protein: Structure, function, and antiviral discovery. Antiviral Res. 207, 105423 (2022).

10. Nguyen, T. H. et al. Targeting west nile virus replication by xanthine inhibitors. Med. Chem. Res. 34, 2558–2582 (2025).

11. Beasley, D. W. C. et al. Envelope protein glycosylation status influences mouse neuroinvasion phenotype of genetic lineage 1 West Nile virus strains. J. Virol. 79, 8339– 8347 (2005).

12. Xie, Xuping et al. Inhibition of Dengue Virus by Targeting Viral NS4B Protein. J. Virol. 85, 11183–11195 (2011).

13. Wang, Y. et al. Mechanistic insights into dengue virus inhibition by a clinical trial compound NITD-688. Proc. Natl. Acad. Sci. U. S. A. 122, e2426922122 (2025).

14. Kiemel, D. et al. Pan-serotype dengue virus inhibitor JNJ-A07 targets NS4A-2K-NS4B interaction with NS2B/NS3 and blocks replication organelle formation. Nat. Commun. 15, 6080 (2024).

15. Miller, S., Sparacio, S. & Bartenschlager, R. Subcellular localization and membrane topology of the Dengue virus type 2 Non-structural protein 4B. J. Biol. Chem. 281, 8854– 8863 (2006).

16. Li, Y. et al. Secondary Structure and Membrane Topology of the Full-Length Dengue Virus NS4B in Micelles. Angew. Chem. Int. Ed Engl. 55, 12068–12072 (2016).

17. Li, Y. et al. Secondary structure and membrane topology of dengue virus NS4B N-terminal 125 amino acids. Biochim. Biophys. Acta 1848, 3150–3157 (2015).

18. Eberhardt, J., Santos-Martins, D., Tillack, A. F. & Forli, S. AutoDock Vina 1.2.0: New Docking Methods, Expanded Force Field, and Python Bindings. J. Chem. Inf. Model. 61, 3891–3898 (2021).

19. Trott, O. & Olson, A. J. AutoDock Vina: improving the speed and accuracy of docking with a new scoring function, efficient optimization, and multithreading. J. Comput. Chem. 31, 455–461 (2010).

20. Schake, P., Bolz, S. N., Linnemann, K. & Schroeder, M. PLIP 2025: introducing protein-protein interactions to the protein-ligand interaction profiler. Nucleic Acids Res. 53, W463–W465 (2025).

21. Lee Ng, M. Ultrastructural Studies of Kunjin Virus-infected Aedes albopictus Cells. J. Gen. Virol. 68, 577–582 (1987).

22. Welsch, S. et al. Composition and Three-Dimensional Architecture of the Dengue Virus Replication and Assembly Sites. Cell Host Microbe 5, 365–375 (2009).

23. Chatel-Chaix, L. et al. A Combined Genetic-Proteomic Approach Identifies Residues within Dengue Virus NS4B Critical for Interaction with NS3 and Viral Replication. J. Virol. 89, 7170–7186 (2015).

24. Bhardwaj, T., Kumar, P. & Giri, R. Investigating the conformational dynamics of Zika virus NS4B protein. Virology 575, 20–35 (2022).

25. Porter, S. S., Gilchrist, T. M., Schrodel, S. & Tai, A. W. Dengue and Zika virus NS4B proteins differ in topology and in determinants of ER membrane protein complex dependency. J. Virol. 99, e0144324 (2025).

26. Miller, S., Kastner, S., Krijnse-Locker, J., Bühler, S. & Bartenschlager, R. The Non-structural Protein 4A of Dengue Virus Is an Integral Membrane Protein Inducing Membrane Alterations in a 2K-regulated Manner*. J. Biol. Chem. 282, 8873–8882 (2007).

27. Lin, D. L. et al. The ER Membrane Protein Complex Promotes Biogenesis of Dengue and Zika Virus Non-structural Multi-pass Transmembrane Proteins to Support Infection. Cell Rep. 27, 1666–1674.e4 (2019).

28. Ngo, A. M. et al. The ER membrane protein complex is required to ensure correct topology and stable expression of flavivirus polyproteins. eLife 8, e48469 (2019).

29. Sali, T. M. et al. Characterization of a Novel Human-Specific STING Agonist that Elicits Antiviral Activity Against Emerging Alphaviruses. PLoS Pathog. 11, e1005324 (2015).

30. Lanciotti, R. S., Lambert, A. J., Holodniy, M., Saavedra, S. & Signor, L. D. C. C. Phylogeny of Zika Virus in Western Hemisphere, 2015. Emerg. Infect. Dis. 22, 933–935 (2016).

31. Smith, J. L. & Hirsch, A. J. Analysis of Serum Anti-Zika Virus Antibodies by Focus Reduction Neutralization Test. Methods Mol. Biol. Clifton NJ 2142, 73–80 (2020).

32. Briese, T., Glass, W. G. & Lipkin, W. I. Detection of West Nile virus sequences in cerebrospinal fluid. Lancet Lond. Engl. 355, 1614–1615 (2000).

33. Kim, G. et al. Easy and accurate protein structure prediction using ColabFold. Nat. Protoc. 20, 620–642 (2025).

34. Jumper, J. et al. Highly accurate protein structure prediction with AlphaFold. Nature 596, 583–589 (2021).

